# High-affinity agonist binding to C5aR results from a cooperative two-site binding mechanism

**DOI:** 10.1101/2020.04.03.024018

**Authors:** Andra C. Dumitru, R. N. V. Krishna Deepak, Heng Liu, Melanie Koehler, Cheng Zhang, Hao Fan, David Alsteens

**Affiliations:** Université catholique de Louvain, Louvain Institute of Biomolecular Science and Technology, 1348 Louvain-la-Neuve, Belgium; Bioinformatics Institute (BII), Agency for Science, Technology and Research (A*STAR), Singapore; Department of Pharmacology and Chemical Biology, School of Medicine, University of Pittsburgh, Pittsburgh, PA15261, USA

**Keywords:** Single-molecule, Atomic force microscopy, FD-based AFM, G-protein-coupled receptors, C5aR, molecular dynamics, ligand-receptor, cooperativity

## Abstract

A current challenge in the field of life sciences is to decipher, in their native environment, the functional activation of cell surface receptors upon binding of complex ligands. Lack of suitable nanoscopic methods has hampered our ability to meet this challenge in an experimental manner. Here, we use for the first time the interplay between atomic force microscopy, steered molecular dynamics and functional assays to elucidate the complex ligand-binding mechanism of C5a with the human G protein-coupled C5a receptor (C5aR). We have identified two independent binding sites acting in concert where the N-terminal C5aR serves as kinetic trap and the transmembrane domain as functional site. Our results corroborate the two-site binding model and clearly identify a cooperative effect between two binding sites within the C5aR. We anticipate that our methodology could be used for development and design of new therapeutic agents to negatively modulate C5aR activity.

## Introduction

The complement C5a anaphylatoxin elicits a variety of immunological responses *in vivo* (Guo and Ward, 2005), such as the stimulated production of pro-inflammatory cytokine by binding to its cognate cell surface receptor, the G-protein-coupled receptor C5a anaphylatoxin chemotactic receptor 1 (C5aR). This interaction has been a topic of interest in the last couple of decades due to its relevance in several inflammatory pathologies, such as asthma, arthritis, sepsis and more recently Alzheimer’s disease and cancer (Ward, 2004; Woodruff, Nandakumar *et al.*, 2011; Klos, Wende *et al.*, 2013). However, the binding mechanism of the C5a ligand to C5aR remains poorly understood at the molecular level hampering the development of new therapeutic agents (Monk, Scola *et al.*, 2007; Klos, Wende *et al.*, 2013). A two-site binding mechanism has been suggested, with the C5a rigid core interacting with both the N-terminus and the second extracellular loop of the receptor (called binding site (BS)) (Siciliano, Rollins *et al.*, 1994), and the C5a flexible C-terminal fragment interacting with the cavity formed by the seven transmembrane (7-TM) helices and involved in the functional activation of C5aR (called effector site (ES)). On one hand, the main interactions at the BS occur between C5aR sulfonated tyrosine residues (Y11 and Y14) and C5a residues R40, R37 and possibly H15 (Siciliano, Rollins *et al.*, 1994; Farzan, Schnitzler *et al.*, 2001; Huber-Lang, Sarma *et al.*, 2003). On the other hand, the C5a R74 is pointed out as a critical residue in the binding to the ES (Siciliano, Rollins *et al.*, 1994; Huber-Lang, Sarma *et al.*, 2003). However, given the absence of the structure of the C5a-C5aR complex, these interactions have never been directly confirmed. In addition, clear and direct evidence of how these two binding sites could act in concert is missing. Understanding this process is likely to illuminate the binding paradigm common to members of the GPCR family that bind large macromolecular ligands.

Although high-resolution structures of GPCRs have emerged during the last decades, these structures are just a snapshot of a specific stabilized state frozen in time and space, among a myriad of possible dynamic conformations. Investigating complex binding mechanisms, in particular to receptors having multiple intramolecular binding sites, remains a remarkably difficult task to date. The dynamic nature of these processes requires the development of new methodologies that have the ability to simultaneously quantify and structurally map ligand-receptor interactions at the single-molecule level in a dynamic setting. Atomic force microscopy (AFM) combines high-resolution imaging from the micrometer to the (sub-)nanometer range with the capability to apply and measure wide dynamic force ranges from pico-to nanoNewtons, which is necessary to characterize the broad spectrum of chemical interactions occurring in living systems. Force-distance curve-based atomic force microscopy (FD-based AFM) has established itself as a powerful method for imaging a variety of biological systems at sub-nanometer resolution and to simultaneously extract quantitative parameters such as topography, adhesion, elasticity and stiffness (Dufrene, Martinez-Martin *et al.*, 2013; Pfreundschuh, Alsteens *et al.*, 2015; Rigato, Rico *et al.*, 2015; Dumitru, Conrard *et al.*, 2018; Dumitru, Poncin *et al.*, 2018). Functionalization of the AFM tip with specific ligands allows specific detection of (bio-)chemical interactions, while simultaneously extracting their structural, thermodynamic and kinetic parameters. Here, we use FD-based AFM to unravel the binding mechanism at the two C5aR sites and to decipher roles of these sites in the kinetic and energetic properties of the binding process. For the first time, using our affinity imaging method, we have successfully extracted binding properties of a single ligand at a molecular sub-site level. The combination of AFM with steered molecular dynamics (SMD) simulation and functional assays revealed new perspectives on the role of allosteric interactions from a kinetic, thermodynamic and functional point of view.

## Results

### C5aR adopts random orientation after reconstitution in lipid membrane

Before characterizing the binding properties of C5a to C5aR, we imaged purified C5aR reconstituted in liposomes and adsorbed on freshly cleaved mica in buffer solution using FD-based AFM. While membrane receptors embedded in their native cellular membrane always have a unique orientation, they lose this original orientation through the reconstitution steps into liposomes (Pfreundschuh, Alsteens *et al.*, 2015). Within lipid bilayers, embedded receptors can adopt both orientations, having either their extracellular or intracellular side exposed to the AFM tip (**Figure 1A**).

**Figure 1.**
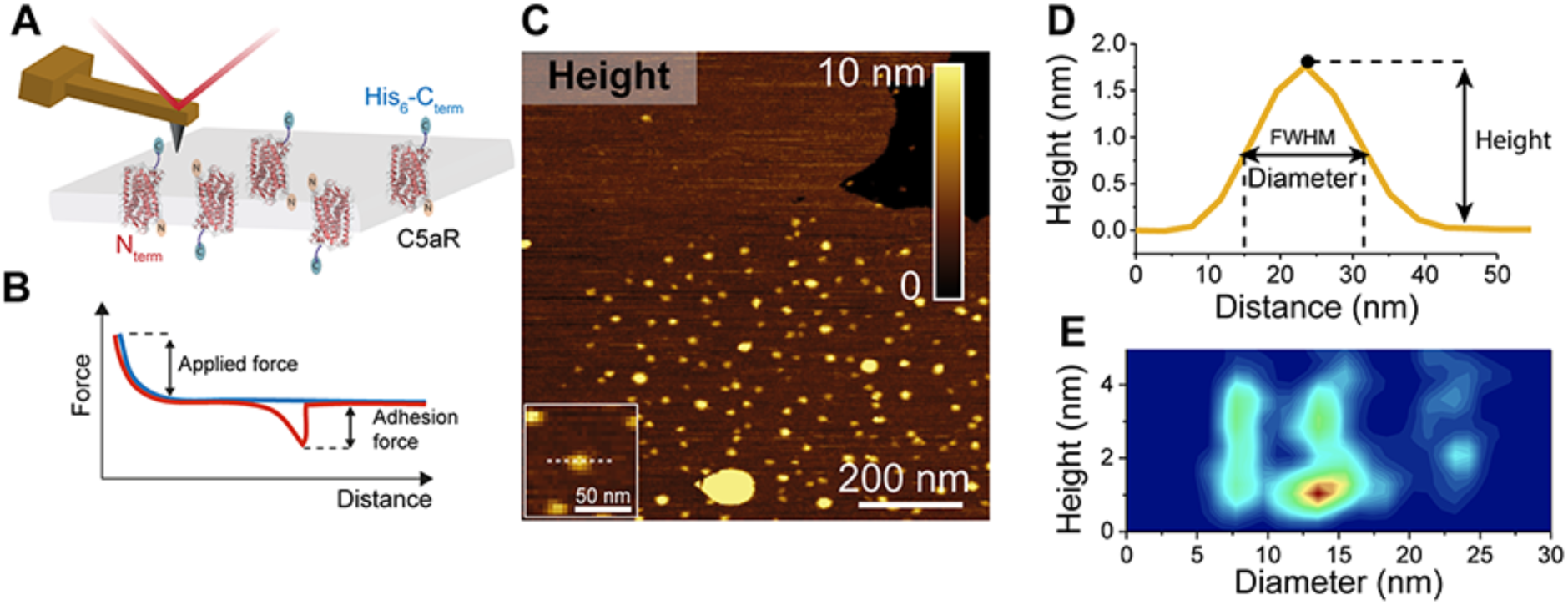
FD-based AFM mapping of C5aR receptors and probing their orientation within the lipid bilayer. **(A)** Orientation of lipid bilayer-embedded C5aR is random: they can adopt two orientations, with either the intracellular C-terminal His_6_-tag or the extracellular N-terminal side facing up. **(B)** In FD-based AFM, the force acting on the cantilever and the distance travelled by the AFM tip are monitored and transformed into a force-distance curve. The applied force is used as feedback during the measurement and the adhesion force is measured as the minimum force in the retraction cycle. **(C)** Overview AFM topography image (height map) of C5aR reconstituted in liposomes and adsorbed on freshly cleaved mica. Sparsely distributed C5aR particles can be observed protruding from the liposomes. Inset: expanded view of a single C5aR particle. The image was acquired with a bare AFM tip. **(D)** Cross-section (white dashed line in **C**) showing a C5aR particle protruding 1.7 nm from the lipid bilayer having a diameter of 16 nm. The diameter was measured as full width at half maximum (FWHM). **(E)** 2D-histogram of height and diameter of C5aR receptors imaged in **C**. The diameter distribution shows three main populations, while the height distribution shows two main peaks. Data in **C** and **E** are representative of at least five independent experiments.

To detect the C5aR orientation within the lipid bilayer, we first imaged our sample at high-resolution using FD-based AFM to possibly discriminate topographical characteristics corresponding to one of the two orientations. In FD-based AFM, an oscillating AFM cantilever is continuously approached and retracted from the sample surface in a sinusoidal manner (**Figure 1A**) and for each pixel of the image a force-distance (FD) curve is recorded (**Figure 1B**). The sample topography along with the adhesion can be simultaneously extracted from each FD curve (**Figure 1B**) (Dufrene, Martinez-Martin *et al.*, 2013; Alsteens, Pfreundschuh *et al.*, 2015; Dumitru, Poncin *et al.*, 2018). The sample imaged by AFM revealed sparsely distributed C5aR particles protruding away from the lipid bilayer surface (**Figure 1C**). The height of the emergent part of C5aR above the lipid bilayer (protrusion height) as well as its diameter (calculated as the full width at half maximum) were extracted for each particle and plotted in a two-dimensional histogram (2D) (**Figures 1D and E**). A bimodal distribution is observed for the heights, with two peaks centered respectively at 1.1 ± 0.5 nm and 2.8 ± 1.2 nm, while the presence of three main populations can be observed for the diameters, with peaks centered at 8.6 ± 0.4 nm (monomers), 14.1 ± 0.3 nm (dimers) and 23.4 ± 1.2 nm (oligomers) (László, Miklós *et al.*, 2008; Alsteens, Pfreundschuh *et al.*, 2015; Liu, Kim *et al.*, 2018). The two peaks of height distribution could represent the heights of the extracellular and the intracellular regions of C5aR protruding from the DOPC/CHS membrane. For the monomers, we clearly observed that both orientations are present without preference (similar density in the 2D-histogram).

### Identification of C5aR orientation using affinity imaging

After having evidenced that embedded receptors show different protrusion height from the membrane, we combined topography and affinity imaging using functionalized AFM tips to specifically identify C5aR intracellular and extracellular sides (**Figure 2**). Silicon tips were functionalized with a poly(ethylene glycol) linker (PEG), followed by grafting of *tris*-*N*-nitrilotriacetic acid (*tris*-NTA) molecules. The individual tetradentate NTA ligand forms a hexagonal complex with Ni^2+^ ions, leaving two remaining binding sites accessible to electron donor nitrogen atoms from the histidine sidechains of the His_8_-tag engineered to the terminal end of a polypeptide. To specifically probe one side of the C5aR, we either used the *tris*-NTA-Ni^2+^ tip to target the intracellular side using the His_8_-tag present at the C5aR C-terminal end (**Figures 2A-D**), or we further derivatized the *tris*-NTA-Ni^2+^ tip with the endogenous C5a ligand to specifically probe the interaction with the ligand-binding site at the C5aR extracellular side (**Figures 2E-H**). Adhesive events were considered to be specific if they were detected at tip-sample distances of 10 ± 5 nm, corresponding to the length of the extended PEG linker, and when the adhesion force was at least two times higher than the noise level (measured at the baseline of the retraction curve, see Methods). Additionally, each specific adhesion event was validated by fitting the extension profile of the PEG linker using the worm-like chain model (Sulchek, Friddle *et al.*, 2006). Representative FD curves are presented in Figures 2D and H showing either unspecific/no interaction FD curves (curves 1-2) or specific adhesion events (curve 3). Control experiments using bare tips or an amino-derivatized tip show either no interaction or unspecific adhesion events (**Figure S1A-D**). Finally, blocking experiments using free C5a in solution or injection of EDTA significantly reduces the binding probability, confirming the specificity of both probed interactions (**Figure S1E-H**).

**Figure 2.**
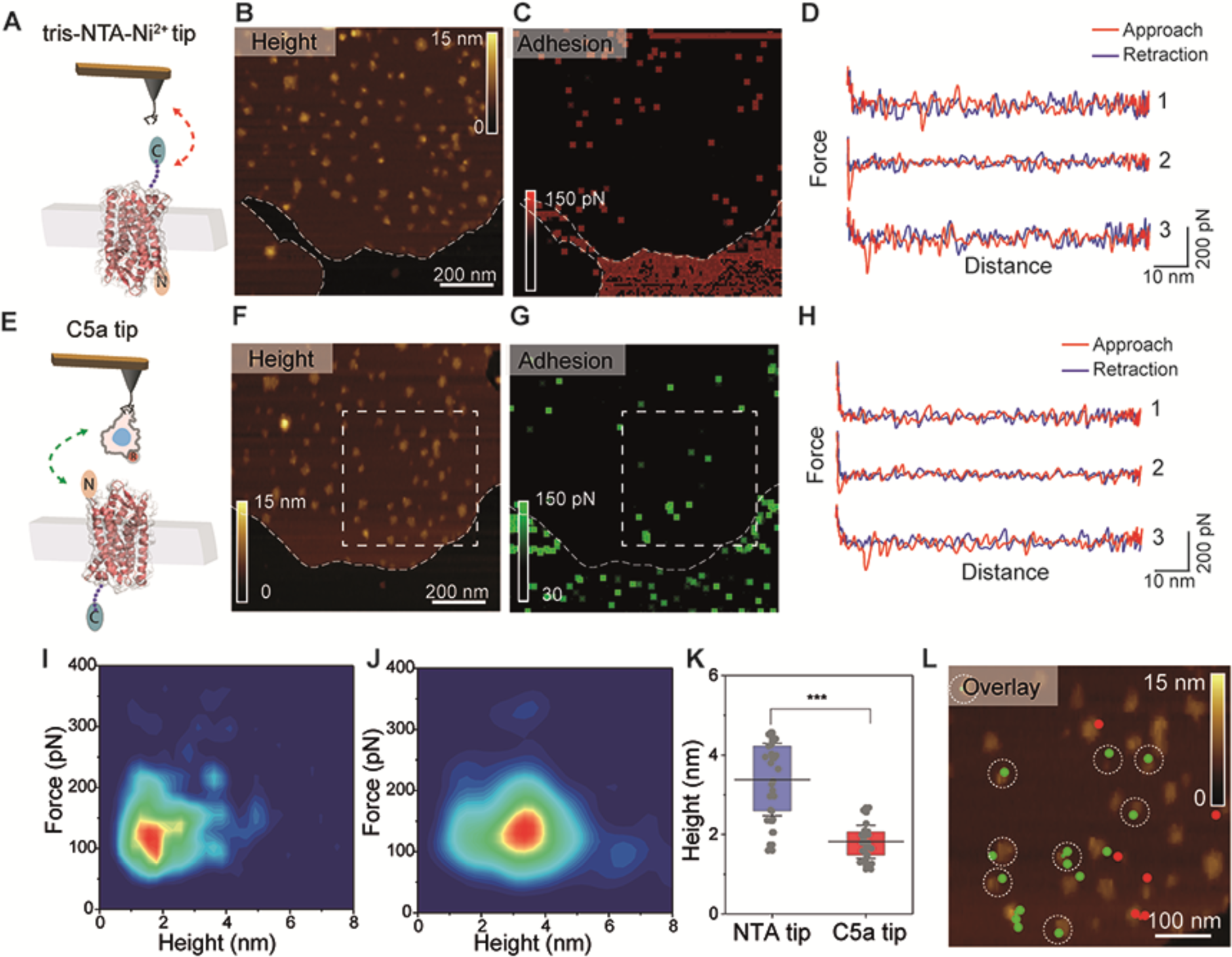
Multiplex probing of C5aR intra- and extracellular binding sites as a method discriminate C5aR orientation within lipid membrane. (**A, E**) Schematic representation of the multiplex experimental setup. Two different AFM tip chemistries were used to target either the His_6_-tag C-terminal end of C5aR using *tris*-NTA-Ni^2+^ functionalized AFM tips (A-D) or the N-terminal end of C5aR using the endogenous C5a ligand (E-H). (**B, C, F, G**) AFM topography and adhesion images were recorded over the same lipid patch with a *tris*-NTA-Ni^2+^ tip (B and C) and a C5a ligand tip (F and H). (**D, H**) Representative FD curves showing either no/unspecific interactions (curves 1 and 2) or specific adhesion events (curve 3) were extracted from the adhesion maps in (**C**) and (**G**) and displayed in (**D**) and (**H**), respectively. (**I, J**) 2D-histograms of force *vs* height for C5a modified tips (**I**) and tris-NTA-Ni^2+^ tips (**J**).(**K**) Height distribution of the receptors interacting with the *tris*-NTA-Ni^2+^ or the C5a tip. Two populations can be clearly distinguished, one below 1.75 nm in height, where C5a tips mostly interact with the extracellular side of C5aR, and another one above 3.5 nm in height, where NTA-Ni^2+^ functionalized tips interact with the intracellular side of C5aR. **(L)** Overlay of the height map region marked by the white square in F and the corresponding specific adhesion events extracted from the same areas in the maps in C and G. Adhesion events between the C5a ligand and the N-terminal side of C5aR are shown as green dots, while the events rising from the tris-NTA-Ni^2+^ AFM tip interaction with the His_6_-tagged C-terminal side of C5aR are displayed as red dots. White circles mark receptors with a height less than 1.75 nm, highlighting that C5a ligand only interact with the N-terminal side of. The overlay image shows how the orientation of single C5aR particles can be identified using our multiplex probing method. Data are representative of at least three independent experiments. Data in K is displayed as mean ± S.D. and the ANOVA OneWay Tukey test was used to report the statistical significance: *** p<0.001. Data are representative of three independent experiments.

For the two types of tip functionalization, the FD curves showing specific adhesion events were analyzed and the interaction force was extracted as well as the height of the C5aR on which the FD curve was recorded. These values were displayed in the form of 2D-histograms of force as a function of height (**Figures 2I and J**). We observed that *tris*-NTA-Ni^2+^ functionalized tips mostly interact with receptors having a protruding height centered at 3.1 ± 1.0 nm together with forces of 125 ± 50 pN, while C5a tips were found to interact specifically with receptors more buried into the membrane (height of 1.7 ± 0.5 nm) and interacting with forces of 110 ± 45 pN. Together, these results confirm that the receptor orientation can be determined using only their protrusion heights as those are significantly different (**Figure 2K**). An overlay of the AFM topography and the specific adhesion events recorded (colored pixels) on the same area with both functionalized tips (C5a tip in red and *tris*-NTA-Ni^2+^ tip in green) reveals the identity of the side exposed to the tip (**Figure 2L**). Receptors having a protrusion height under a threshold of 1.75 nm were encircled. Together, these data confirm the possibility to identify with a high-probability (> 95%) the receptors oriented in their native state. Therefore, this height criterion will be used in the following force spectroscopy experiments to validate our measurements and to only probe native state receptors.

### C5a is a high-affinity agonist of C5aR

Next, we wanted to characterize C5a ligand binding to C5aR (**Figure 3A**). To this end, we simultaneously recorded FD-based AFM height images and adhesion maps (**Figures 3B and C**) and extracted FD curves located on C5aR having their native orientation (based on our height criterion) (**Figure 3D**). Generally, force-probing methods such as FD-based AFM measure the strength of single bonds under an externally applied force. Described first by the Bell-Evans model (Evans and Ritchie, 1997), an external force stressing a bond reduces the activation-energy barrier toward dissociation and, hence, reduces the lifetime (τ) of the ligand-receptor pair (**Figure 3E**). The model predicts that far from equilibrium, the rupture force (e.g., binding strength) of the ligand-receptor bond is proportional to the logarithm of the loading rate (LR), which describes the force applied over time. Recently, Friddle, Noy and de Yoreo (FNdY) introduced a model to interpret the nonlinearity of the rupture forces measured over a wide range of LRs and suggested that this nonlinearity arises through the re-formation of bonds at small LRs (Friddle, Noy *et al.*, 2012). This model provides direct access to the equilibrium free energy ΔG_bu_ between bound and unbound states (see Methods). The non-linear oscillating approach and retraction movement of the AFM tip with respect to the sample results in a wide variety of velocities explored during the rupture of the bonds established between the ligand derivatized tip and C5aR (**Figure 3F**). To further increase the range of velocities explored, we combined the force-volume (FV) mode at low speed to reach low LRs and FD-based AFM to explore unbinding behavior at high LRs (**Figure 3F**). For each FD showing a specific adhesion event, we extracted the binding force and the LR measured as the slope of the force *versus* time curve just before the rupture (**Figure 3G**). When plotting the resulting binding forces as a function of the LRs (also called dynamic force spectroscopy (DFS) plot) on a semi-logarithmic scale (**Figure 3H**), a non-linear dependency of the force with the loading rate is observed as predicted by the FNdY model. Using this model, we extracted the equilibrium force F_eq_, as well as thermodynamic and kinetic parameters such as the binding equilibrium free energy ΔG_bu_ and the receptor-ligand half-life τ_0.5_. An equilibrium force F_eq_ of 46 ± 7 pN, a binding equilibrium free energy ΔG_bu_ of -13.6 ± 4.1 kcal mol^-1^ and a dissociation constant K_d_ of ≈ 6.09 nM were found. The dissociation constant was calculated using the relation ΔG_bu_ = k_B_T x ln(0.018 K_d_) with 0.018 l mol^-1^ being the partial molar volume of water. The K_d_ estimated using our single-molecule method is the same order of magnitude as previously reported values, ranging from 1-10 nM, based on radioactive ligand binding assays (Gerber, Meng *et al.*, 2001; Robertson, Rappas *et al.*, 2018). These results highlight that FD-based AFM was suitable to quantify the kinetic and thermodynamic binding of a large ligand interacting with its receptor via multiple orthosteric sites.

**Figure 3.**
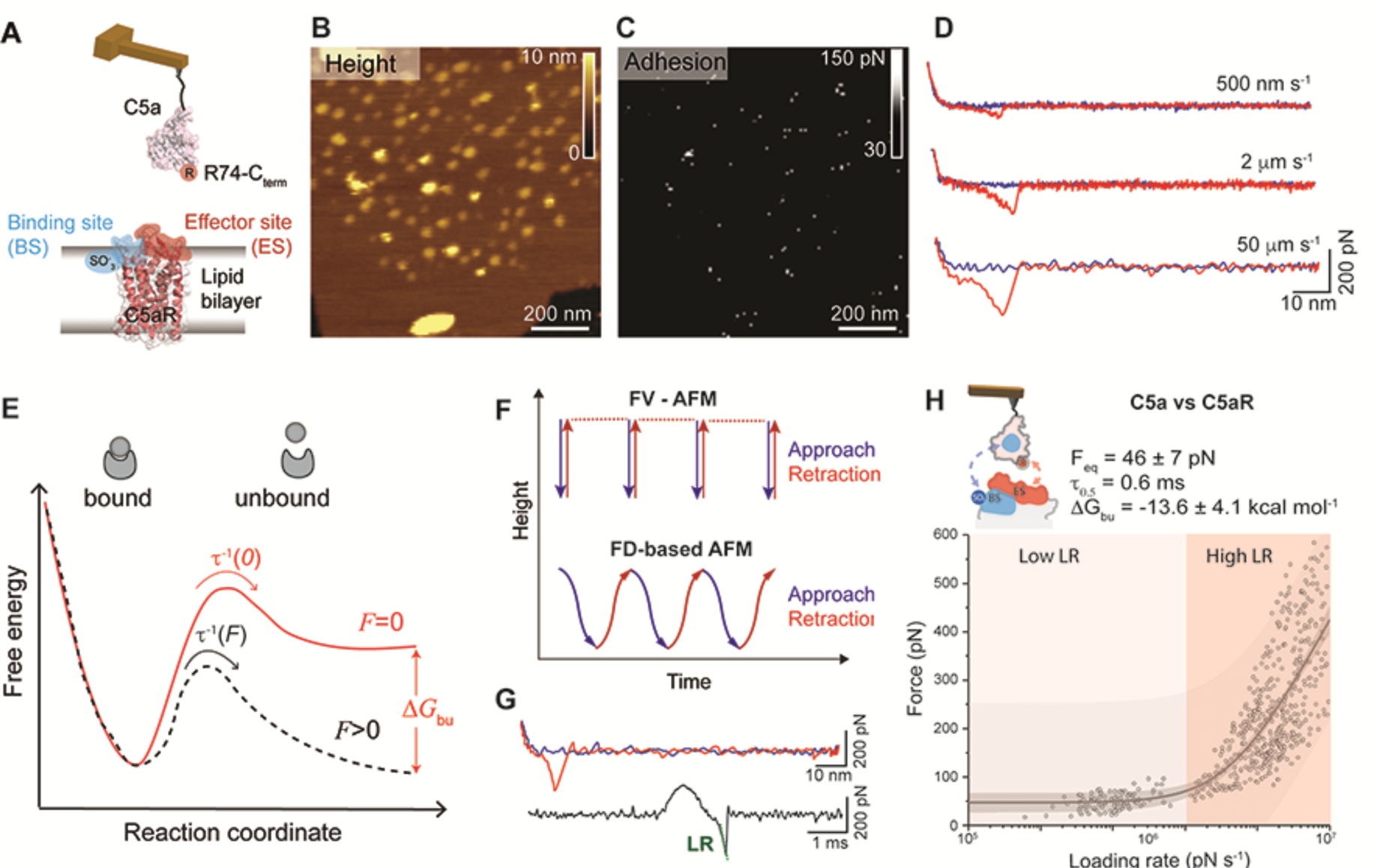
Probing the kinetic and thermodynamic parameters underlying C5a ligand binding to C5aR. **(A)** Two-site binding model of the C5a-C5aR interaction: the binding site (BS) at its N-terminus (shaded in blue) and the functionally important effector site (ES) at the extracellular region (shaded in red). Sulfonated residues (SO_3_^-^) at the BS and Arg (R) residue at the C-terminal of the C5a ligand are thought to play a key role in stabilizing the interaction. (**B**) Height image and adhesion maps (**C**) recorded while probing C5aR embedded in the lipid bilayer with a C5a modified AFM tip. **(D)** The interaction between C5a and C5aR was probed over a wide range of LRs by variating the retraction speed in the force-distance curves. Low LRs were explored at 500 nm s^-1^ and 2 µm s^-1^ pulling speeds, while high LRs were reached at 50 µm s^-1^ pulling speed.**(E)** Extracting the parameters describing the C5a-C5aR free energy landscape. A ligand-receptor bond can be described using a simple two-state model, where the bound state resides in an energy valley and is separated from the unbound state by an energy barrier. The transition state must be overcome to separate ligand and receptor. τ^-1^(F) and τ^-1^(0) are residence times linked to the transition rates for crossing the energy barrier under an applied force F and at zero force, respectively. Δ*G*_bu_ is the free-energy difference between bound and unbound state. **(F)** Force-volume (FV)-AFM and FD-based AFM were used to explore low LRs and high LRs, respectively. For each pixel of the topography the tip is approached and retracted using a linear (FV-AFM) or oscillating movement (FD-based AFM). **(G)** A force-distance curve can be displayed as a force-time curve, from which the loading rate can be extracted via the slope of the curve just before bond rupture. **(H)** DFS plot showing the loading rate-dependent interaction forces of the C5a ligand with C5aR. Data combines rupture forces obtained at lower LRs (10^2^-10^5^ pN.s^-1^) and higher LRs (10^5^-10^7^ pN.s^-1^) Fitting the data using the Friddle–Noy–de Yoreo model (thin grey line) provides average *F*_eq_, Δ*G*_bu_ and residence time (τ_0.5_) values with errors representing the s.e.m. Each circle represents one measurement. Darker shaded areas represent 99% confidence intervals, and lighter shaded areas represent 99% of prediction intervals. For each condition, data are representative of at least three independent experiments.

### MD simulations identify key residues involved in antagonist binding to C5aR

Although it is thought that C5a binds C5aR through two orthosteric sites, a crystal structure of the C5a-C5aR complex is missing. To better investigate the key residues responsible for the high affinity interaction, we turned our attention to PMX53, a well-known C5aR full antagonist that mimics the structure of the C-terminal segment of C5a (March, Proctor *et al.*, 2004; Monk, Scola *et al.*, 2007; Woodruff, Nandakumar *et al.*, 2011). The C5aR-PMX53 crystal structure (Liu, Kim *et al.*, 2018) has been recently elucidated and we can hypothesize that given their high structural similarities, the cyclic hexapeptide PMX53 and the C-terminal segment interact with C5aR through similar residues. To get more insight into the PMX53-C5aR binding dynamics we performed molecular dynamics (MD) and steered MD (SMD) simulations (**Figure 4**). Over the course of the 300 ns unrestrained MD simulation, the C5aR structure remained stable with no significant structural changes as evidenced from the root-mean-square deviation (RMSD) profile (**Figure S2A**). Apart from the N- and C-terminal regions of C5aR, most of the structural fluctuations were observed primarily in the second extracellular loop (ECL2) region. The termini of TM5 and TM7 with the 7-TM core of the protein remained fairly stable (**Figure S2B**). No major structural rearrangements were observed with respect to the PMX53 molecule, which remained tightly bound to C5aR, forming several hydrogen bonds (**Figure S3A and B**). Key intermolecular interactions between C5aR and PMX53 present in the initial crystal structure such as the D282-R6_PMX53_ salt-bridge remained stable throughout the entire 300 ns (**Figure 4**). The Y258-R6_PMX53_ cation-π interaction was broken (R6_PMX53_ CZ and Y258 ring-centroid distance > 6.0 Å) halfway through the simulation but the two residues remained close to each other (**Figure 4G**). Disruption of the cation-π interaction allowed R6_PMX53_ to interact with D282 in a head-on manner (**Figure 4G and Figure S3C**). W5_PMX53_ and R6_PMX53_ saddled Y258 but did not interact directly during the production run (**Figure 4G and Figure S3C**).

**Figure 4.**
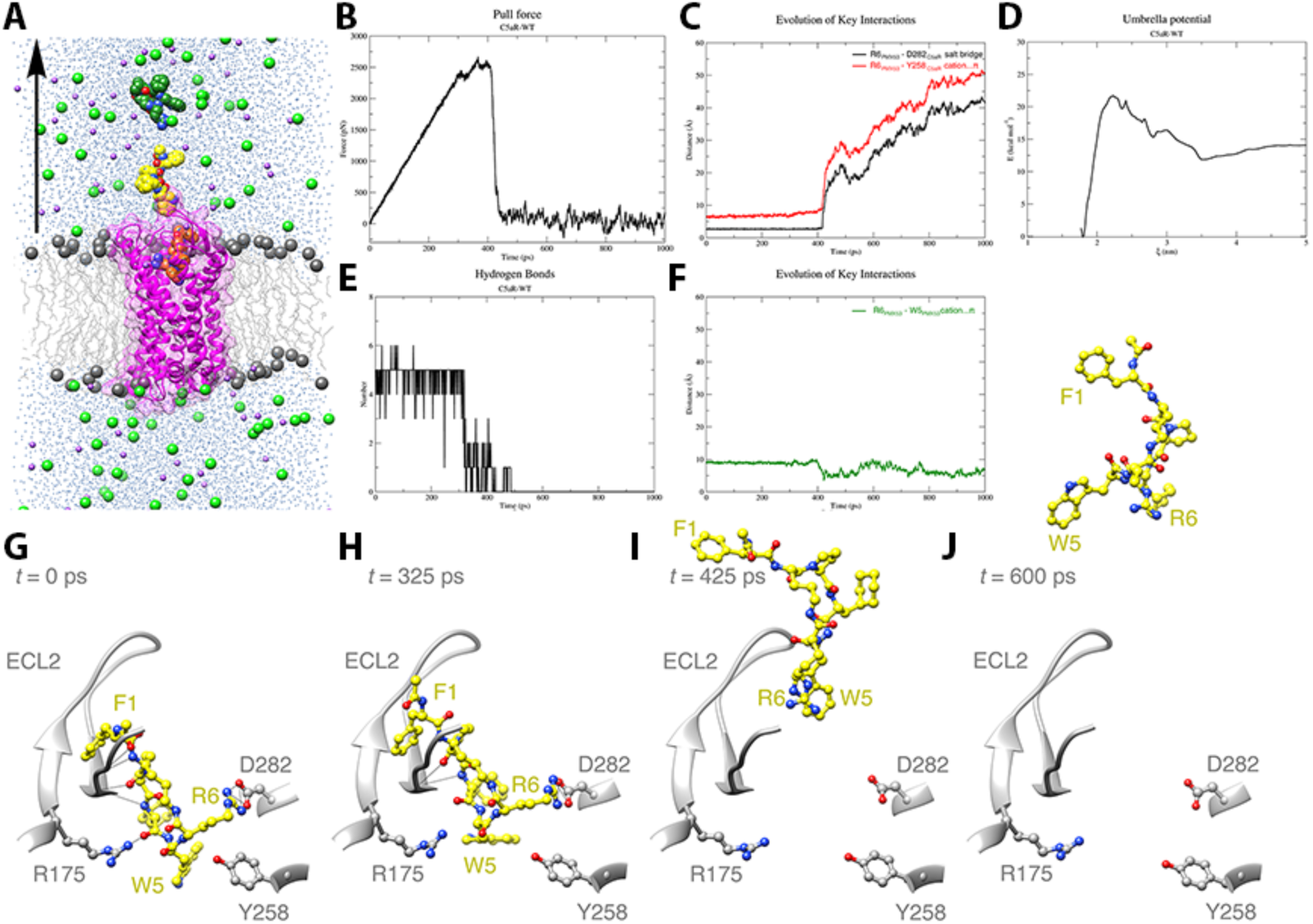
Steered Molecular Dynamics (SMD) or Center-of-mass (COM) pulling simulation of C5aR (WT) – PMX53 complex. **(A)** Cut-through section of C5aR (WT)-PMX53-POPC system used for equilibrium MD and steered MD simulations. C5aR is shown in ribbon representation (magenta), embedded in a POPC bilayer (gray) with the headgroup phosphorous atoms shown in sphere representation and the rest of the lipid molecule shown in wire representation. TIP3P water molecules are colored blue, Na+ ions purple and Cl-ions green. Conformations of PMX53 at t = 0 ps, t = 500 ps, and t = 1000 ps derived from the COM pulling simulation are shown in orange, yellow, and dark green colors, respectively. The black arrow is along the z-axis and indicates the direction of pulling of PMX53. **(B)** Plot showing force (pN) vs time (ps) profile obtained for the C5aR (WT)-PMX53 system with a pulling rate of 5 nm/ns. **(C)** Plot showing the number of intermolecular hydrogen bonds (H bonds) formed/broken between the ECL2 region (residues 174-196) of C5aR (WT) and PMX53 over the course of the pulling simulation. **(D)** Evolution of key intermolecular interactions between C5aR (WT) and PMX53, namely the R6PMX53-D282C5aR salt-bridge (black), and the R6PMX53-Y258C5aR cation…π interaction (red) over the course of the pulling simulation. **(E)** Evolution of key intramolecular interaction R6PMX53-W5PMX53 cation…π interaction (green) in PMX53 over the course of the pulling simulation. **(F)** Potential of mean force profile calculated for the dissociation of PMX53 from C5aR (WT) using WHAM analysis following umbrella sampling simulations for the C5aR (WT)-PMX53 system. The average PMF profile calculated using bootstrap analysis is presented in the Supplementary Figure. XX. **(G)** Position and conformation of PMX53 at t = 0 ps during pulling simulation (pull force = 7.62×10-5 pN) where R6PMX53 stably and directly interacts with D282C5aR as compared to the conformation observed in the starting crystal structure conformation. In this conformation, PMX53 forms extensive H bond interactions (shown as black lines) with the residues of C5aR (WT), especially with residues of ECL2. (H) Position and conformation of PMX53 at t = 325 ps during pulling simulation (pull force = 2386.14 pN) where key non-covalent interactions between PMX53 and C5aR (WT) begin to break and R6PMX53 and W5PMX53 are being pulled away from Y258C5aR and D282C5aR. A number of HBonds between PMX53 and ECL2 also as broken or are in the process of being broken under the influence of the applied force.**(I)** Position and conformation of PMX53 at t = 425 ps during pulling simulation (pull force = 879.08 pN) where the PMX53 molecule has been pulled further away with the R6PMX53-D282C5aR salt-bridge and the R6PMX53-Y258C5aR cation…π interaction being completely broken. **(J)** Position and conformation of PMX53 at t = 600 ps during pulling simulation (pull force = 29.48 pN) where the ligand is completely unbound from the receptor.

### PMX53 dissociates from C5aR in two critical steps

Steered molecular dynamics has been successfully employed for studying biological phenomenon such as stability of α-amyloid protofibrils (Lemkul and Bevan, 2010), substrate translocation by membrane transporters (Shi, Quick *et al.*, 2008), and interaction of GPCR ligands with their cognate receptors (Yuan, Raniolo *et al.*, 2018). We employed SMD or center-of-mass (COM) pulling simulations on the final configuration of the 300 ns equilibrium simulation to gain an atomistic insight into the molecular events that occur during the dissociation of PMX53 from the C5aR binding pocket. Akin to AFM experiments, in the pulling simulations, the bound PMX53 molecule was pulled away from the C5aR binding pocket by applying an external force along the *z*-axis (**Figures 4A and B and Movie 1**). The force *vs.* time profile of the pulling simulation is presented in Figure 4B. The application of force on the PMX53 molecule led to a gradual build-up of force until a critical point was reached that was sufficient to break the key intermolecular interactions to allow the dissociation of the bound molecule.

The plot showed two such critical points, a minor drop in force around *t* = 308 ps and a major drop in force around *t* = 425 ps. After the major drop, the PMX53 molecule was mostly unbound. We analyzed the evolution of various intermolecular non-covalent interactions between C5aR and PMX53 during the pulling simulation. The analysis revealed that shortly after *t* = 308 ps time-point numerous hydrogen bonds, almost half of which were formed between the ligand and the residues of the C5aR ECL2 region, were broken, resulting in a brief drop in force (**Figures 4C and H**). Further, the critical D282-R6_PMX53_ salt-bridge and the Y258-R6_PMX53_ cation-π were completely broken around *t* = 425 ps time-point when the pulling force was maximal (**Figures 4D and I**). Following the breakage of these critical interactions, the PMX53 molecule adopted a more compact conformation facilitated by the formation of an intramolecular R6_PMX53_-W5_PMX53_ cation-π interaction (**Figures 4E and J**).

We also performed umbrella sampling simulations on the configurations generated from the non-equilibrium SMD trajectories to calculate the free energy profile of the PMX53 binding/dissociation events. The weighted histogram analysis method (WHAM) was employed to obtain the potential of mean force (PMF) curve from which the *Δ*G of PMX53 binding was deduced (**Figure 4F**). Bootstrap analysis was used to estimate the statistical errors, and the average PMF profile along with the corresponding standard deviation values are plotted in Figure S4. On the basis of the PMF profile, we obtained a *Δ*G_bu_ value of -13.8 kcal mol^-1^.

### PMX53 binds to C5aR with a high affinity

In parallel with the pulling simulations, we quantified by FD-based AFM the free-energy landscape of the PMX53-C5aR interaction and the role of the key residues identified by our simulation. To this end, we tethered the high-affinity PMX53 antagonist to the AFM tip and then measured its interactions with C5aR (**Figure S5A**) and two C5aR mutants identified through MD simulation (**Figure S5B**,**C**). Fitting the experimental data with the FNdY model provided an equilibrium force F_eq_ of 50 ± 9 pN corresponding to a binding equilibrium free energy ΔG_bu_ of - 13.7 ± 4.9 kcal mol^-1^ for PMX53-C5aR interaction. This value is very similar to the value determined by the SMD simulation for the PMX53-C5aR. The affinity is also similar in magnitude to the value determined for the C5a ligand although the latter, due to its large size, interacts on multiple binding sites on the receptor. This observation is probably a result of the PMX53 reduced size and the various substitutions (compared to the C5a native C-terminus) that makes it accommodate better within the binding pocket. The PMX53 affinity constant K_d_ of 4.7 nM is in good agreement with previous studies where values between 1-50 nM were found depending on the species and cell type (Woodruff, Strachan *et al.*, 2001; Seow, Lim *et al.*, 2016).

### C5aR^R175/Y258^ influences the binding kinetics

Next, we tested C5aR mutations within the ES, as pointed by the above MD and SMD simulations. We designed two C5aR mutants (D282A and R175V/Y258V) located in the ES and mediating direct polar interactions (R175 and D282) or water-mediated polar interactions (R175). We performed several functional assays with the C5aR and the two mutants and observed a strong reduction on the G_i_-protein activation in response to increased concentration of C5a (**Figure S5**), confirming the crucial role played by these residues in the modulation of C5aR’s functional state. We then probed the interaction by FD-based AFM with the PMX53 on both mutants. Thermodynamic analysis using the FNdY model only revealed a slight reduction of the *Δ*G_bu_ from -13.7 ± 4.9 kcal mol^-1^ for the C5aR^WT^ to -13.3 ± 1.0 kcal mol^-1^ for the C5aR^R175V/Y258V^ double mutant (**Figure S5B**). This slight reduction in ΔG_bu_ has also been confirmed by MD and SMD simulations using the PMX53-C5aR^R175V/Y258V^ double mutant system following the same protocol used for the PMX53-C5aR^WT^ system. The most striking observation from the MD simulation of PMX53 with the C5aR double-mutant was the reduction in the number of hydrogen bonds formed between C5aR^R175V/Y258V^ and PMX53, particularly involving residues from ECL2. The R175V mutation causes a 66% reduction in the number of hydrogen bonds formed between ECL2 and PMX53 (1.57 ± 0.91) as compared to the wild-type (4.62 ± 0.93; Figure S4E). The R6_PMX53_-D282 salt-bridge remained stable throughout the 300ns production run whereas the Y258V mutation caused a change in the stability of the W5_PMX53_ orientation (**Figure S4D**). When PMX53 was pulled away from the double-mutant C5aR using a similar SMD protocol, we observed a marked drop in the force required for the ligand to dissociate (**Figure S4B**). The inter- and intramolecular non-covalent interactions behave in a similar fashion as the wild-type but break much earlier (**Figures S4C-E**). The hydrogen bonds between ECL2 and PMX53 broke much earlier around *t* = 200 ps (**Figure S4E**) as compared to the wild-type (*t* = 308 ps) while the R6_PMX53_-D282 salt-bridge breaks shortly thereafter, but earlier than the wild-type (**Figure S4D**). The PMF profile for the double-mutant shows a significant drop (∼43%) in the height of the energy barrier crossed during PMX53 dissociation (−12.2 kcal mol^-1^ for double-mutant vs. -21.5 kcal mol^-1^ for WT) although resulting in a slight reduction (∼12%) in ΔG_bu_ (−12.2 kcal mol^-1^ for double-mutant vs. -13.8 kcal mol^-1^ for WT) (**Figure S4F**). Results from our MD and SMD studies are in good agreement with our experimental data obtained by AFM where we observed a slight decrease in the ΔG_bu_ (∼ 3%) but a much important reduction in residence time (∼ 40%) that can be directly linked with the reduction of the height of the energy barrier crossed during PMX53 dissociation. Altogether, these results suggest that the R175 and Y258 residues play a key role in the kinetics of the interaction while at the same time being less crucial for the thermodynamics.

### C5aR^D282^ is critical for binding thermodynamics

Finally, we studied the PMX53 binding to the C5aR^D282A^ mutant by FD-based AFM (**Figure S5C**). The analysis of the DFS plot with the FNdY model revealed a strong reduction of the free energy (∼ 35%) leading to a ΔG_bu of_ -7.7 ± 1.9 kcal mol^-1^, while the residence time remains unchanged (0.1 ms). MD and SMD simulations were attempted on this mutant, with no success in obtaining convincing results for the umbrella sampling simulations due to largely reduced PMX53-C5aR^D282A^ interactions. Yet, both experimental and simulation experiments suggest a strong reduction of the interactions due to the single point mutation in the receptor, thus underlying the important role of D282 in the stabilization of the PMX53 into the binding pocket.

### C5a C-terminus weakly binds C5aR effector site

As PMX53 is a peptide that mimics the structure of the C-terminal segment of C5a, we wondered whether the C-terminal segment of C5a could also bind to the C5aR ES with high-affinity. To prevent the interaction between the core of C5a and the sulfation sites at the BS, the C5aRΔTyr mutant with mutations Y11F and Y14F sites was generated (Farzan, Schnitzler *et al.*, 2001). The lack of sulfation on the C5aRΔTyr mutant was then validated by Western Blot using anti-sulfated tyrosine antibodies (**Figure 5A**). The interaction between the C5a ligand and C5aRΔTyr was measured and the dependence of the rupture force with the loading rate was plotted in the DFS graph in Figure 5B. A non-linear dependency of the rupture force with the loading rate was again observed and the FNdY model was used to fit the data. We extracted an equilibrium force F_eq_ of 32 ± 14 pN, a binding equilibrium free energy ΔG_bu_ of -4.7 ± 3.4 kcal mol^-1^ and a receptor-ligand half-residence time, τ_0.5_ of 0.5 ms. The calculated ΔG_bu_ corresponds to a dissociation constant K_d_ of ≈ 20 mM and corresponds to a surprisingly low-affinity, in contrast to the high-affinity binding of C5aR^WT^. To further increase the accuracy of the extracted parameters, we also recorded specific binding events at lower LRs by oscillating the cantilever at frequencies of 1-10 Hz (**Figure 5C**). The measured forces over the low LRs regime (10^2^-10^5^ pN.s^-1^) align well with binding forces obtained at higher LRs (10^5^-10^7^ pN.s^-1^). The superimposition of the rupture forces obtained in the lower LRs regime (10^2^-10^5^ pN.s^-1^) to the high LRs makes it possible to better visualize the existence of a force plateau in the close-to-equilibrium regime as predicted by the FNdY model (Friddle, Noy *et al.*, 2012). The fit of the whole data set (low and high LRs) gives very close values for the extracted parameters (**Figures 5B and C**).

**Figure 5.**
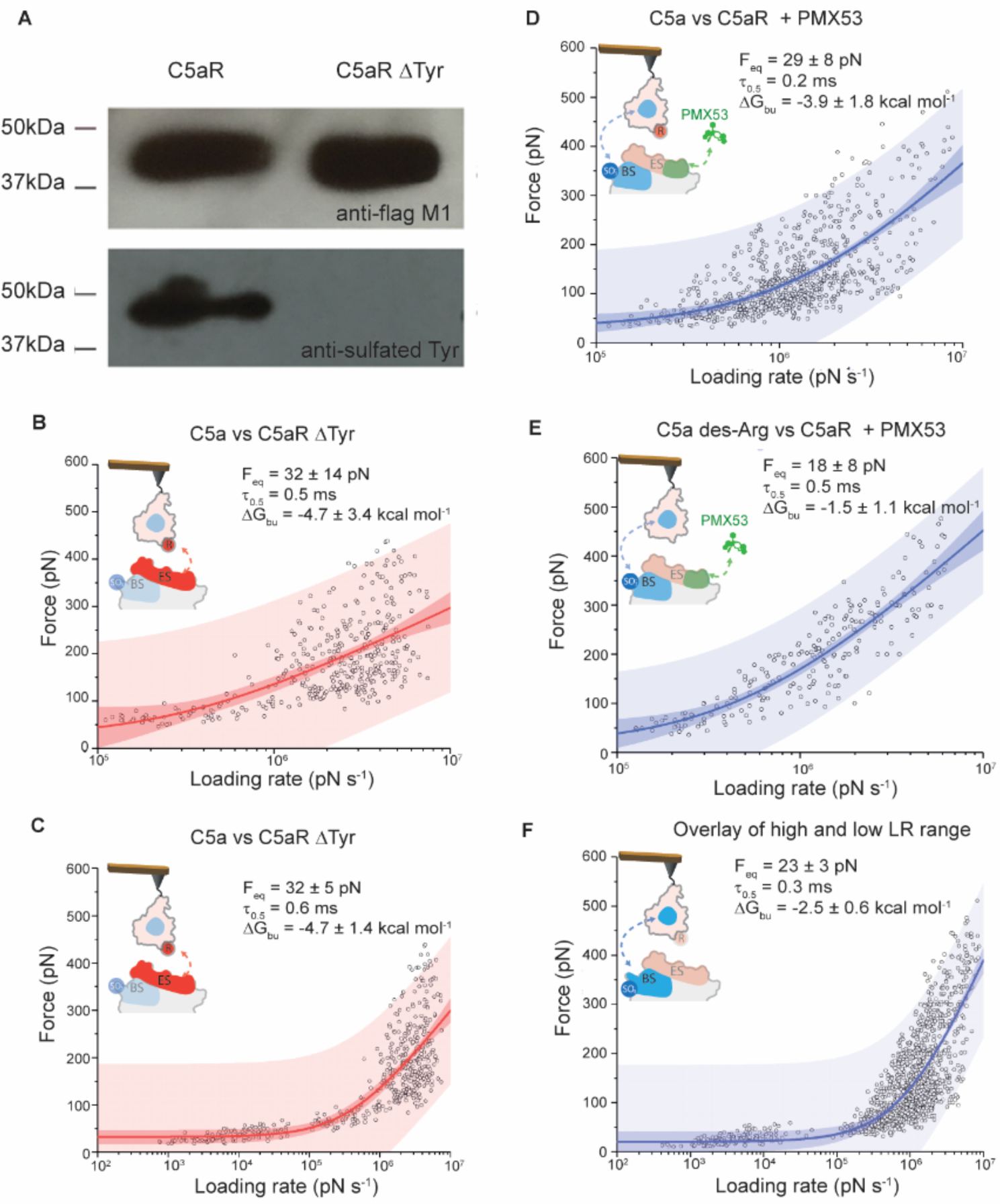
Probing the kinetic and thermodynamic parameters underlying C5a ligand binding to C5aR at the sub-site level. **(A)** Western blot analysis of C5aR sulfonation. Detection of sulfonation of wt C5aR and C5aRΔTyr with Y11F and F14F mutations (C5aR-d2Y). Both constructs are with an N-terminal FLAG tag, which was detected by an anti-Flag M1 antibody. The two mutations in C5aRΔTyr eliminated the sulfonation on C5aR. For each condition, data are representative of at least three independent experiments. **(B**,**C)**. Exploring C5a binding to the effector site of C5aR. In **(B)** DFS plot showing the loading rate-dependent interaction forces of the C5a ligand with the C5aR missing a Tyr residue at the N-terminal end. The DFS plot in **(C)** combines rupture forces obtained at lower and higher LRs (10^2^-10^7^ pN.s^-1^), covering the close-to-equilibrium and far-from-equilibrium binding strengths. **(D**,**E**,**F)** Probing the interactions of the C5a ligand with the C5aR binding site. DFS plots showing the loading rate-dependent interaction forces of: **(D)** the C5a ligand with C5aR complexed in presence of the PMX53 antagonist, **(E)** C5a des-ArgC5a des-Arg with C5aR complexed with the PMX53 antagonist and **(F)** DFS plot combining rupture forces obtained at lower and higher LRs (10^2^-10^7^ pN.s^-1^), covering the close-to-equilibrium and far-from-equilibrium binding strengths. In panels (B,C,D,E,F) fitting the data using the Friddle–Noy–de Yoreo model (thin lines) provides average *F*_eq_, Δ*G*_bu_ and residence time (τ_0.5_) values with errors representing the s.e.m. Each circle represents one measurement. Darker shaded areas represent 99% confidence intervals, and lighter shaded areas represent 99% of prediction intervals. For each condition, data are representative of at least three independent experiments.

### The rigid core of C5a has a low affinity for the C5aR binding site

As the binding of C5a to the C5aR ES alone fails to explain the high-affinity interaction, we also explored the binding between the C5a rigid core and the C5aR BS (**Figures 5D-F**). To specifically target the BS, we abolished the interactions that could be established with the ES site using two different approaches: (*i*) injection of free PMX53 (**Figure 5D**) and (*ii*) using C5a des-Arg + PMX53 (**Figure 5E**). C5a des-Arg is an endogenous truncated C5a derivative lacking the R74 at its C-terminal end, which is known to be crucial for the interaction with the ES (Cain, Coughlan *et al.*, 2001; Higginbottom, Cain *et al.*, 2005).

Fitting the DFS plots with the FNdY model revealed that the inner core of the C5a binds to C5aR BS with free-energy values (ΔG_bu_) of -3.9 kcal mol^-1^ and -1.5 kcal mol^-1^, depending on the presence or absence of the R74 residue. The most energetically unfavored complex was observed when the C5a des-Arg ligand was probed along with PMX53 injection (ΔG_bu_ ≈ -1.5 kcal mol^-1^). We also looked into the kinetics aspect of the C5a-BS interaction and quantified the complex stability in terms of residence time. τ_0.5_ values of 0.2 and 0.5 ms were obtained for the two conditions (**Figures 5D,E**). The most stable complex was the one formed by the C5a des-Arg ligand probed in the presence of free PMX53 in solution (τ_0.5_ ≈ 0.5 ms). Since the exact structural details of C5a interaction with its receptor have not been revealed experimentally yet, it is difficult to engineer specific mutants that would completely abrogate the interaction at the ES. As, in both cases the interactions to the ES are abolished by PMX53, the high-affinity antagonist, we only probed the interaction to the BS site and combine the results obtained above (**Figure 5F**). Fitting with the FNDY model gave an equilibrium force F_eq_ of 23 ± 9 pN and a binding equilibrium free energy ΔG_bu_ of -2.5 ± 1.9 kcal mol^-1^. The calculated ΔG_bu_ corresponds to a very low dissociation constant K_d_ of ≈ 0.8 M. To further increase the accuracy of the extracted parameters, we also explored the close-to-equilibrium regime of the binding of C5a to the C5aR BS (**Figure 5F**). We obtained very close F_eq_ values (23 ± 3 pN for the whole set *vs* 23 ± 9 pN for only the high LR range). The residence time and binding free-energy also remain unchanged.

The two-fold difference in ΔG_bu_ between C5a binding at the ES and the BS, together with the higher affinity for the ES, suggest that the interactions at the ES largely dominate the binding of C5a to C5aR, while still weak when measured individually. Nevertheless, we hypothesize the binding to multiple intramolecular sites, such as the ES and BS, stabilizes the overall binding, increasing the ligand residence time and therefore acting as a kinetic trap with the purpose of raising the local binding concentration.

### High-affinity C5a binding results from positive allostery between ES and BS

In the light of the results obtained above, we further analyzed the functional activation of the C5aR^WT^ and C5aR^D282A^ mutant exposed to C5a and C5a des-Arg (**Figures 6A,B and S5**). We observed for the C5aR^D282A^ mutant a strong reduction on the G_i_-protein activation in response to increased concentration of C5a (**Figure 6B**), revealing that in addition to its important role in the ligand affinity, this residue also modulates C5aR functional state. Otherwise, functional assays performed with the truncated C5a des-Arg suggested that the loss of the interactions between C5aR-D282 and C5a-R74 only slightly affect the normal C5aR function (**Figure 6A**).

**Figure 6.**
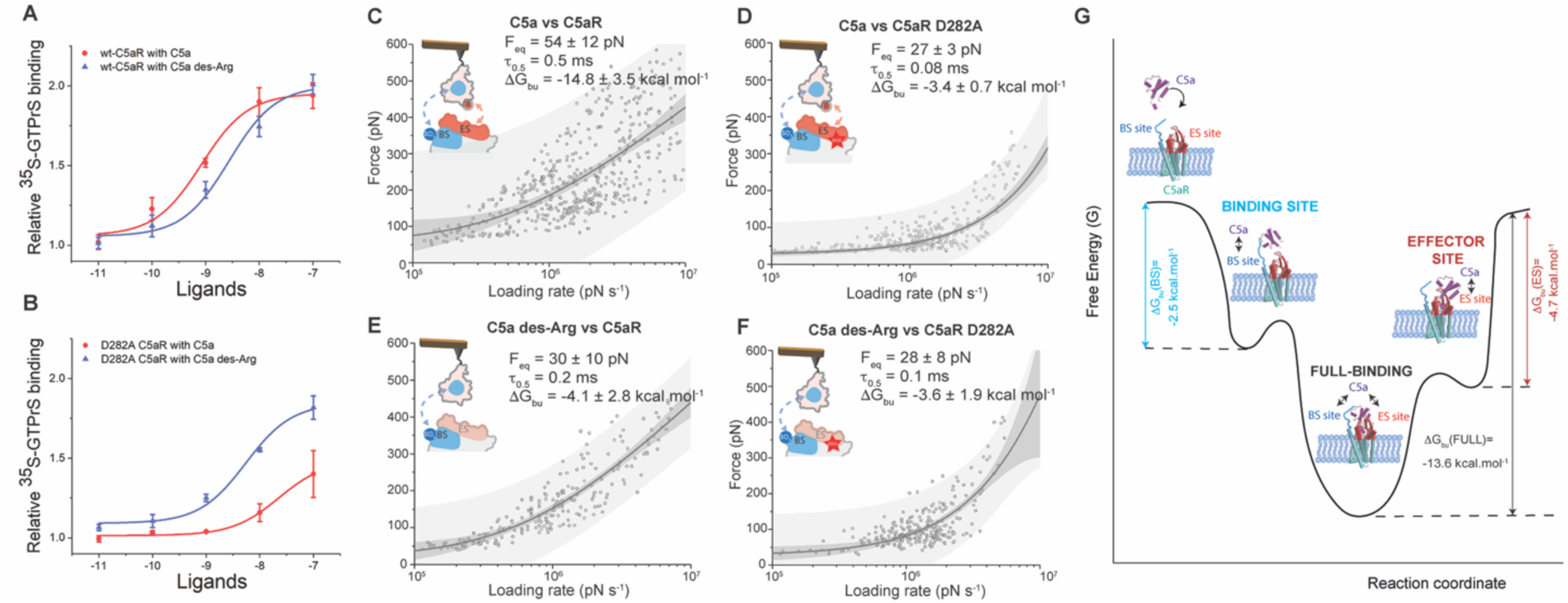
Probing the kinetic and thermodynamic parameters underlying ligand binding to C5aR mutants. (**A, B**) Dose-response curves of C5a and C5a des-Arg in activating G_i_ protein through the action on the wtC5aR and C5aR D282A mutant. The activation of G_i_ protein was determined by measuring the binding of ^35^S-GTPγS to Gi. DFS plots showing the loading rate-dependent interaction forces of the C5a ligand with the C5aR (**C**), C5a ligand with the C5aR D282A (**D**), C5a des-Arg interacting with C5aR (**E**) and C5a des-Arg interacting with C5aR D282A (**F**). Fitting the data using the Friddle–Noy–de Yoreo model (thin lines) provides average *F*_eq_, Δ*G*_bu_ and residence time (τ_0.5_) values with errors representing the s.e.m. Each circle represents one measurement. Darker shaded areas represent 99% confidence intervals, and lighter shaded areas represent 99% of prediction intervals. For each condition, data are representative of at least three independent experiments. (**G**) Cooperative binding of C5a to C5aR though a two-site binding mechanism. Illustration of the free-energy binding landscape of C5a binding to C5aR. Δ*G*_bu_ gives the free-energy difference between the ligand-bound and unbound states and is indicated for each binding site (BS, ES and BS+ES) by vertical arrows. A positive allosteric interaction is measured when both binding sites (BS and ES) are occupied, as revealed by a significantly higher Δ*G*_bu_ for the full binding of the C5a.

Next, we wanted to know to what extend the binding of the C5a ligand to the ES influences the binding in the overall C5a-C5aR complex (**Figures 6C-F**). We tested the influence of the C5a-R74 residue (using C5a des-Arg mutant) and the C5aR-D282 mutation. For the C5a-C5aR interaction (probed in the high LRs range) we obtain very similar results as previously determined (**Figure 3**): an equilibrium force F_eq_ of 54 ± 12 pN, a binding equilibrium free energy ΔG_bu_ of -14.8 ± 3.5 kcal mol^-1^ and a dissociation constant K_d_ of ≈1 nM. Deletion of the R74 residue in the C terminal segment or mutation of the D282 residue in the C5aR lead to strong effect with a three-fold drop of the ΔG_bu_, in the low-affinity regime (**Figures 6D-F**). These results corroborated our previous observation that each sub-site (ES or BS) taken individually interacts with low-affinity in the molar or millimolar range, alluding to a possible cooperative action between both sites in the overall binding mechanism. To address this, we performed a two-sample t-test distribution with the hypothesis that the ΔG_bu_ of the full ligand is larger than the sum of the ΔG_bu_ of each binding sites (ΔG_bu_ (BS+ES)). However, although it is still difficult or impossible to develop general models for multiple bonds within a molecular assembly, we can nevertheless assume that a maximum ΔG_bu_ would be observed for bonds failing cooperatively when loaded in parallel, that would mean ΔG_bu_ (BS+ES)= ΔG_bu_ (BS)+ ΔG_bu_ (ES)= -7.2 ± 2.0 kcal mol^-1^. The t-test confirms (p-value=0.964) that the full-ligand binding free-energy (ΔG_bu_(C5a)) is significantly higher than the sum of the binding free-energy of both sites measured individually (ΔG_bu_ (BS+ES)), confirming a positive allosteric interactions between the two orthosteric binding sites, establishing the full interaction with the C5a. Our AFM experiments further suggest that the interaction between C5a-R74 and C5aR-D282 plays a pivotal role into this cooperative mechanism. Indeed, the single point mutation into the C5aR (D282A) or truncated C5a des-Arg is sufficient to completely abolish this high affinity interaction state (**Figure 6D-F**).

## Discussion

GPCRs represent the largest human membrane protein family, having overall more than 800 members, and constitute a “control panel” of the cell (Latorraca, Venkatakrishnan *et al.*, 2017). As predominant actors in cells, GPCRs are intensively studied as drug targets, where in particular C5aR has long been suggested as a new promising anti-inflammatory target. Intensive research on C5aR has led to the design of several antagonists including the peptide antagonist PMX53 and several non-peptide antagonists such as NDT9513727 and avacopan. PMX53 is a potent orthosteric antagonist with insurmountable action (Seow, Lim *et al.*, 2016), although its peptidic nature has limited its clinical development (Klos, Wende *et al.*, 2013). Among the current available non-peptide antagonists, only avacopan showed sufficient therapeutic efficacy to advance into late-stage clinical trials (Bekker, Dairaghi *et al.*, 2016; Jayne, Bruchfeld *et al.*, 2017). Recent structural studies on C5aR revealed that the non-peptide antagonists (including avacopan and NDT9513727) are actually allosteric modulators with highly reversible action (Liu, Kim *et al.*, 2018; Robertson, Rappas *et al.*, 2018). Further development of orthosteric non-peptide antagonists could be preferred as they may exhibit an insurmountable action similar to PMX53. Corroborated with previous studies revealing the structural basis for the action of PMX53 (Liu, Kim *et al.*, 2018), our kinetic and thermodynamic insights of PMX53 binding to the receptor effector site confirmed by MD and SMD simulations, shed more light into the activation mechanism of C5aR and the amino acid residues involved, which could be useful for future drug discovery studies.

Atomic-resolution structures are now available for more than 50 different GPCRs and over 250 of their complexes with different ligands (Shimada, Ueda *et al.*, 2018). Crystal structures of C5aR in complex with NDT9513727, PMX53 and avacopan have recently been reported (Liu, Kim *et al.*, 2018; Robertson, Rappas *et al.*, 2018). However, despite the dramatic progress during the last decade in deciphering the structural insights of C5aR activation mechanism, none of those recent structural studies have been performed with the C5a ligand. In addition, the function of GPCRs depends critically on their ability to change shape, transitioning among distinct conformations, while crystal structures only depict discrete snapshots of a dynamic process. Although for some GPCRs several small drug candidates have been developed using solely structure-based drug design methodologies (Rodríguez, Ranganathan *et al.*, 2015), a full understanding of the dynamic properties of GPCRs is preferred and probably essential for future drug development, especially for those with large peptide or protein ligands. Here, we introduced an FD-based AFM approach and a new experimental strategy to extract the kinetic and thermodynamic parameters governing large-ligand binding to multiple orthosteric binding sites of receptors in physiologically relevant conditions. We also used MD and SMD simulations as a powerful complementary method to our experimental approach, allowing us to gain new insights into the binding pocket structure and the important residues involved in the specific recognition of ligands.

Our study addressed the complex binding process of a large ligand to a GPCR. C5a, a 74-aa glycoprotein binds to C5aR through two distinct and physically separated binding sites, namely the effector and binding site (Siciliano, Rollins *et al.*, 1994). While the existence of the two-site binding motif has been previously reported (Siciliano, Rollins *et al.*, 1994), the functional relationship between the two sites was missing until now. Our method enabled, for the first time, to probe multiple ligand binding sites at the sub-site level in order to study their respective contribution to the overall binding. We demonstrated that both orthosteric ligand binding sites interact with the ligand with a low affinity when working independently. Interestingly, when acting in a concerted manner, the interaction rises into a high-affinity interaction, suggesting a cooperativity between both orthosteric binding sites (**Figure 6G**). This cooperativity effect resulting from multiple binding site is supported by the theory that predicts (Williams, 2003; Sulchek, Friddle *et al.*, 2006) an enormous increase of the time scale needed for ligand dissociation upon cooperative binding.

Through our experimental approach combining AFM and simulations, we were able to capture the “cryptic” binding pockets of C5a into C5aR and to reconstruct the binding free-energy landscape for this complex binding mechanism. The importance of the D282 at the extracellular face of TM7 was also put in evidence. Although already predicted to form an important interaction with R74 of C5a, this interaction remained so far a mystery since some D282 mutants were showed to be relatively unresponsive to C5a but sensitive to C5a des-Arg and analogs (Cain, Coughlan *et al.*, 2001; Cain, Higginbottom *et al.*, 2003). More recently, it has been shown that the truncated C5a des-Arg bind C5a in an entirely different orientation, which could be an intermediate state. For the first time, we decipher that this interaction is only established for the full C5a ligand and that the ligand-binding in its high-affinity state involved the concerted action of both the binding and effector sites. We envision that this better understanding of the dynamic binding of C5a to C5aR in physiologically relevant conditions will open new avenues in the rational design of finely tuned drugs. Ultimately, this approach will serve as a valuable tool to further develop and test agonists and antagonists to other GPCRs with macromolecular ligands.

## Materials and methods

### C5aR^WT^ expression, purification and Western Blot

The wild type C5aR and mutant were expressed in mammalian HEK-293S GnTI^-^ cells (ATCC) using the BacMam method (Dukkipati, Park *et al.*, 2008). All constructs were cloned into a vector engineered from pFastBac (Invitrogen) by introducing a CMV promoter (Dukkipati, Park *et al.*, 2008). All protein was expressed with a C-terminal His_8_ tag and an N-terminal Flag tag. Baculovirus was generated by the Bac-to-Bac method (Invitrogen). The mammalian HEK-293S GnTI^-^ cells were cultured in suspension at 37°C and under 5% CO_2_. The cells were infected at a density of 4×10^6^ ml^-1^ with baculovirus and then harvested after 24h.

To purify the protein, infected cells were lysed in buffer containing 10 mM Tris pH 7.5, 150 μg ml^-1^ benzamidine, 0.2 μg·ml^-1^ leupeptin and 2 mg·ml^-1^ iodoacetamide. The cell membrane was collected by centrifugation at 24,000 g for 40 min at 4°C and then solubilized in buffer containing 20 mM HEPES pH 7.5, 750 mM NaCl, 1% dodecyl maltoside (DDM), 0.2% cholesterol hemisuccinate (CHS), 0.2% sodium cholate, 20% glycerol, 150 μg·ml^-1^ benzamidine, 0.2 μg·ml^-1^ leupeptin, 2 mg·ml^-1^ iodoacetamide and 5 U/l Salt Active Nuclease (Sigma) for 1 h at 4 °C. The supernatant was collected after centrifugation at 24,000 g for 40 min, and incubated with Ni-NTA agarose resin (Clontech) in batch for overnight at 4°C. The resin was washed three times in batch with buffer comprised of 20 mM HEPES pH 7.5, 500 mM NaCl, 0.1% DDM, 0.02% CHS, 150 μg·ml^-1^ benzamidine, 0.2 μg·ml^-1^ leupeptin and 20 mM imidazole, and then transferred to a gravity column. After extensive washing, the protein was eluted in wash buffer with 400 mM imidazole and 2 mM CaCl_2_. The eluted protein was loaded onto anti-Flag M1 antibody affinity resin. After washing with buffer containing 20 mM HEPES pH 7.5, 100 mM NaCl, 0.1% DDM, 0.02% CHS, 2 mM CaCl_2_, the protein was eluted with buffer containing 20 mM HEPES, pH 7.5, 100 mM NaCl, 0.1% DDM, 0.02% CHS, 200 μg·ml^-1^ Flag peptide and 5 mM EDTA. The protein was further purified by size exclusion chromatography with buffer containing 20 mM HEPES pH 7.5, 100 mM NaCl, 0.05% DDM, 0.01% CHS.

Mouse anti-FLAG M1 antibody and mouse anti-Sulfotyrosine antibody (Sigma) were used to detect the purified wild type C5aR and C5aRΔTyr with Y11F and F14F mutations respectively in the western blotting assays.

### C5aR mutants expression, purification and ^35^S-GTPγS binding assay

Mutant variants (D282A, D282N and R175V/Y258V) were generated based on the wtC5aR construct and fully sequenced. Mutant variants were expressed following the same method as for wtC5aR except for some modifications. HEK-293S cells expressing each mutant were pelleted by centrifugation and resuspended in 20 ml buffer containing 20 mM HEPES pH 7.5, 100 mM NaCl, 0.2 μg/ml leupeptin and 150 μg/ml benzamidine. After 20 min incubation at 25 °C, 20 ml 2X solubilization buffer containing 20 mM HEPES pH 7.5, 100 mM NaCl, 1% dodecyl-maltoside (DDM), 0.2% cholesterol hemisuccinate (CHS), 20% glycerol, 0.2 μg/ml leupeptin, 150 μg/ml benzamidine and 5 U Salt Active Nuclease (Sigma) was added. Cell membranes were solubilized for 1.5 hour at 4 °C. The supernatant was collected by centrifugation at 24,000 g for 30 min at 4 °C, and then incubated with anti-Flag M2 antibody affinity resin (Sigma) for 1.5 hour at 4°C. After washing the resin with buffer containing 20 mM HEPES pH 7.5, 100 mM NaCl, 0.1% DDM, 0.02% CHS, 0.2 μg/ml leupeptin, and 150 μg/ml benzamidine, the protein was eluted from M2 resin using the buffer containing 20 mM HEPES pH 7.5, 100 mM NaCl, 0.1% DDM, 0.02% CHS and 200 μg/ml Flag peptide (GL Biochem). The protein was further purified by size exclusion chromatography with the same buffer as for wtC5aR.

For the ^35^S-GTPγS binding assays, the membrane of HEK293S GnTI^-^ cells expressing wtC5aR (∼200 µg/ml) or mutant variants was incubated with 200 nM purified G_i_ protein for 30 minutes on ice in buffer containing 20 mM HEPES pH 7.5, 100 mM NaCl, 5mM MgCl_2_, 3 μg/ml BSA, 0.1μM TCEP, and 5μM GDP to get the receptor and G_i_ complex. Next, 25 μL aliquots of the pre-formed complex were mixed with 225 μL reaction buffer containing 20 mM HEPES, pH 7.5, 100 mM NaCl, 5mM MgCl_2_, 3 μg/ml BSA, 0.1μM TCEP, 1μM GDP, 35 pM ^35^S-GTPγS (Perkin Elmer) and C5a (R&D Systems). After additional 15 min incubation at 25 °C, the reaction was terminated by adding 5 ml of cold wash buffer containing 20 mM HEPES pH 7.5, 100 mM NaCl and 5mM MgCl_2_, and filtering through glass fiber filters (Millipore Sigma). After washing the filters twice with 5 ml cold wash buffer, the filters were incubated with 5 ml of CytoScint liquid scintillation cocktail (MP Biomedicals). The radiation of bound ^35^S-GTPγS was measured on a Beckman LS6500 scintillation counter to determine the binding of ^35^S-GTPγS to G_i_ induced by C5aR activation. The data analysis was performed using GraphPad Prism 6 (GraphPad Software). Results are shown as mean ± s.d. from 3 independent experiments.

### C5aR liposomes preparation

C5aR liposomes were prepared according to previously published method(Pfreundschuh, Alsteens *et al.*, 2015). The empty liposomes were prepared from a mix of DOPC (1,2-Dioleoyl-sn-glycero-3-phosphocholine) (Avanti lipids) and CHS (Sigma). DOPC and CHS were dissolved in chloroform at a 10:1 (wt:wt) ratio, then mixed and dried. The well-mixed DOPC/CHS was re-suspended and dissolved in buffer containing 20 mM HEPES pH 7.5, 100 mM NaCl, 1% octylglucoside (OG) under sonication on ice. Aliquots of dissolved DOPC/CHS lipids were flash-frozen in liquid nitrogen and stored at -80 °C. To reconstitute C5aR in liposomes, protein and lipids were mixed at a 10 µM:1mM final ratio, and incubated on ice for 2h. The detergent was removed by biobeads (Bio-rad) and extensive dialysis.

### C5aR preparation for AFM measurements

The reconstituted C5aR sample solution (either wt-C5aR or mutants) was 20-fold diluted in fusion buffer solution (20 mM HEPES, 300 mM NaCl, 25 mM MgCl_2_) and adsorbed on freshly cleaved mica for 15 minutes. After rinsing with imaging buffer (20 mM HEPES and 300 mM NaCl) the sample was transferred to the AFM.

### Functionalization of AFM tips

Rectangular Si_3_N_4_ AFM cantilevers with silicon tips (BioLever mini, Bruker) were first cleaned with chloroform for 10 min, rinsed with ethanol, N_2_ dried and then cleaned for 15 min in an ultraviolet radiation and ozone cleaner (UV-O, Jetlight, CA, USA). For the aminofunctionalization, the cantilevers were immersed in an ethanolamine solution (3.3 g ethanolamine in 6.6 ml DMSO) overnight and then rinsed in DMSO (3 x 1 min) and ethanol (3 x 1 min), followed by N_2_ drying (Wildling, Unterauer *et al.*, 2011). This was followed by the N-hydroxysuccinimide (NHS)-PEG_27_-acetal linker attachment. A 1 mg portion of the NHS-PEG_27_-acetal linker (JKU, Linz, Austria) was diluted in 0.5 ml chloroform with 30 µl triethylamine and the cantilevers were immersed in this solution for 2 h. After 3 rinsing steps of 10 min in chlorofom and N_2_ drying, the cantilevers were immersed in a 1% citric acid solution for 10 minutes, rinsed with pure water (3 x 5 min) and dryed with N_2_ once more. Tris-NTA-derivatized AFM cantilevers were obtained by pipetting 100 µl of a 100 µM tris-nitrilotriacetic amine trifluoroacetate (Toronto Research Chemicals, Canada) (tris-NTA) solution onto the cantilevers, followed by the addition of 2 μl of a freshly prepared 1 M NaCNBH_3_ solution. The cantilevers were incubated for 1 h, then 5 μl of a 1 M ethanolamine solution pH 8.0 were added for 10 minutes to quench the reaction. Tris-NTA cantilevers were further used to obtain C5a- or C5a des-Arg-derivatized tips. For this purpose, 100 μl of a 1 μM C5a or C5a des-Arg protein solution was premixed with 5 μl NiCl_2_ 5 mM and the mixture was pipetted onto the tris-NTA cantilevers. After 2h of incubation time, the cantileveres were washed in HEPES buffer 3 x 5 minutes.

To functionalize AFM cantilevers with the PMX53 antagonist (Ace-Phe-[{Orn}-Pro-{D-Cha}-Trp-Arg]), aminofunctionalized cantilevers were immersed for 2 h in a solution prepared by mixing 1 mg of NHS-PEG_27_-maleimide(Wildling, Unterauer *et al.*, 2011) (JKU, Linz, Austria) dissolved in 0.5 ml of chloroform with 30 μl of triethylamine, then washed with chloroform and dried with N_2_. The cystein bearing peptide Cys-Gly_3_-Phe-[{Orn}-Pro-{D-Cha}-Trp-Arg] (PMX53-Gly_3_-Cys) was obtained from GL Biochem (Shanghai). A 100 μl solution of Cys-Gly_3_-PMX53 1 mM was premixed with 2 μl of EDTA (100 mM, pH 7.5), 5 μl of HEPES (1 M, pH 7.5), 2 μl of TCEP hydrochloride (100 mM) and 2 μl of HEPES (1 M, pH 9.6), then pipetted over the AFM cantilevers. After 3 h of reaction, cantilevers were washed with PBS 3 x 5 minutes.

### FD-based AFM

AFM experiments were performed with a Multimode 8 AFM equipped with a Nanoscope V controller (Bruker, Santa Barbara, CA, USA) operated in “PeakForce Tapping QNM mode”. All measurements were carried out in imaging buffer at room temperature (≈24°C). For the high-resolution characterization of C5aR topographical features, triangular Si_3_N_4_ cantilevers (Scanasyst-Fluid+, Bruker) with a sharpened tetrahedral silicon tip of ≈2 nm radius, nominal spring constants of 0.35 N/m and resonance frequency in liquid of ≈75 kHz were used. Multiparametric FD-based AFM measurements with derivatized tips were carried out using BioLever mini cantilevers (Bruker, Santa Barbara) having nominal spring constants of 0.1 N/m and resonance frequency in liquid of ≈25 kHz. The spring constant was calibrated at the end of each experiment for all cantilevers used in this study using the thermal noise method(Butt and Jaschke, 1995).

In FD-based AFM measurements, the AFM cantilever is oscillated in a sinusoidal manner well below its resonance frequency, while the sample surface is contoured pixel-by-pixel. For each approach-retraction cycle of the oscillating cantilever, a force-distance curve is recorded. Multiparametric FD-based AFM height, Young’s modulus and adhesion maps are then obtained from a pixel-by-pixel reconstruction of the acquired data. Overview FD-based AFM maps were acquired by scanning the sample at 1 Hz and a resolution of 512 x 512 pixels, using a force setpoint of ≈150 pN, a 2 kHz oscillation frequency and a peak-to-peak oscillation amplitude of 100 nm. Adhesion maps were recorded using a scan rate of 0.2 Hz and 256 x 256 pixels. The functionalized AFM cantilever was oscillated at 0.25 kHz with peak-to-peak oscillation amplitudes of 100 nm. To vertically oscillate the AFM tip at 1–10 Hz, FD-based AFM was conducted in the ramp mode with a force setpoint of 200 pN, an approach velocity of 1 μm.s^-1^, retraction velocities of 0.5-2 μm.s^-1^, a ramp size of 150 nm and no surface delay.

### Control Experiments

Several control experiments were designed to ensure that the measured interactions were indeed specific and the functionalization of the AFM tip successful. Adhesion maps of C5aR reconstituted samples were imaged with unmodified or ethanolamine-coated AFM tips (**Figure S1A, D**). We also tested C5a ligand binding before and after injection of free C5a on the sample surface (**Figure S1C**,**D**). In another approach, tris-NTA binding to C-terminal of C5aR was tested in the presence of 10 mM EDTA (**Figure S1E**,**F**).

### Data analysis

Raw images were analyzed using the Gwyddion 2.5 free software. Force-distance curves were analyzed using the Nanoscope Analysis 1.80 Software (Bruker). Individual force-distance curves corresponding to specific adhesion events were extracted and further analyzed using the OriginLab software. Adhesion forces were calculated as the minimum force in the retraction segment of the force-distance curve and the loading rate was measured as the slope of the force versus time curve just before rupture. The noise level was calculated by doing a linear fit of the retraction part of the force distance curve and calculating the standard deviation. We obtained noise values between 10-15 pN and set a threshold for the specific unbinding events above 25 pN. Dynamic Force Spectroscopy (DFS) graphs were obtained by plotting the loading-rate dependence of the adhesion force and a nonlinear iterative fitting algorithm (Levenberg-Marquardt) was used with the FNdY model to extract kinetic and thermodynamic parameters of the interactions. The fits were plotted along with the 99% confidence intervals and 99% prediction intervals.

The robustness of the FNdY fit was tested for the dataset in Figureure 3H using MATLAB. We maintained one of the three fit parameters (x_β_, F_eq_, k_off_) constant and variated the two others parameters (**Figure S6**). The robustness of a fit is usually evaluated in terms of R^2^ values, but R^2^ is not an optimal choice in a nonlinear regime as the total sum-of-squares (TSS) is not equal to the regression sum-of-squares (REGSS) plus the residual sum-of-squares (RSS), as is the case in linear regression. To circumvent the issue of the low performance of R^2^ and its inappropriateness for nonlinear data analysis, we calculated the sum of the squared differences between the experimental data values and the FNdY fits using the tested parameters. This performance value, *i.e.* the sum of the squared differences, shows how far the data points are from the regression line on average, so low values are indicators of good fit parameters, while high values indicate a poor fit parameter. 2D color maps were reconstructed with color scales showing log-values of the sum of the squared differences between the experimental data values and the FNdY fits with the tested parameters. Minimum values (dark blue) are indicators of the best parameters, while maximum values (dark red) correspond to regions where the fit is poor.

### Molecular Dynamics simulation system setup

The dual antagonist-bound C5aR structure complexed with PMX53 and avacopan in the orthosteric and allosteric sites, respectively, solved by Liu et al. [PDB ID: 6C1R] was used for setup of the simulation systems. All atoms other than those of C5aR and PMX53 (avacopan, solvent, lipids, etc.) were removed along with the engineered N-terminal cytochrome *b*_*262*_ RIL (BRIL). All non-terminal missing regions (234-236, 308-312) were modeled using MODELLER v9.13 *via* the *Model Loops/ Refine Structure* module available in UCSF Chimera (Pettersen, Goddard *et al.*, 2004). A total of 500 structures with the missing loop regions were modeled and the one with the best zDOPE score was selected for preparation of the system for molecular dynamics (MD) simulations.

For the PMX53-C5aR double-mutant system, the R175V and Y258V mutations were introduced into the WT-C5aR-PMX53 system using the *Rotamers* module available in USCF Chimera. The N- and C-termini of C5aR were acetylated and amidated, respectively. The C5aR-PMX53 complex was then embedded in a lipid bilayer comprising 164 POPC (1-palmitoyl-2-oleoyl-*sn-* glycerol-3-phosphocholine) molecules (82 each on the upper and lower layers) using the CHARMM-GUI server. The receptor-antagonist-lipid system was then solvated with 27000 TIP3P water molecules, and NaCl at a concentration of 150 mM was added. The final dimensions of the system were ∼ 79.1 Å x 79.1 Å x 170 Å comprising ∼108,000 atoms. CHARMM36 force field(Klauda, Venable *et al.*, 2010) parameters were used to model protein, lipids, ions, and water molecules. For PMX53, force field parameters were assigned by analogy using CHARMM general force field (CGenFF) *via* the ParamChem server(Ghosh, Marru *et al.*, 2011).

GROMACS v5.1.2 (Ref.(Abraham, Murtola *et al.*, 2015)) was used for performing all the simulations. Short-range non-bonded interactions were calculated with a 1.2 nm cut-off, and the particle mesh Ewald (PME) algorithm(Darden, York *et al.*, 1993) was employed for calculation of long-range electrostatics. LINCS(Hess, Bekker *et al.*, 1997) algorithm was used to constraint all H-atom containing bonds. The system was first energy minimization using steepest decent algorithm. Subsequently, the system was equilibrated in a stepwise manner, first in an NVT ensemble (three steps, 50ps each with 1fs time step) maintained at 310 K by Berendsen coupling. The system was then equilibrated in an NPT ensemble (three steps, 100ps each with 2fs time step) maintained at 310 K and 1.0 bar using Berendsen coupling. The harmonic position restraints applied to the heavy atoms of C5aR, PMX53 and POPC were reduced gradually at each of the six equilibration steps to ensure thorough equilibration of the system. Following equilibration, all restraints were removed and production runs were carried out by maintaining the temperature (310 K) and pressure (1.0 bar) with the help of Nosé-Hoover thermostat and Parrinello-Rahman barostat, respectively. Pressure coupling was carried out semi-isotropically for NPT equilibration and production runs. Finally, a production run of 300 ns was carried out.

### Center-of-Mass (COM) Pulling and Umbrella Sampling Simulations

The resultant configuration of the 300 ns production run was used for performing the COM pulling simulations. The final configuration was equilibrated for 100ps in an NPT ensemble. Subsequently, with positional restraints placed only on the C5aR molecule in the *z-*direction, the bound PMX53 cyclic peptide was pulled away from C5aR binding pocket. The pulling simulation was carried out over 1 ns along the *z*-axis with a pull rate of 0.005 nm ps^-1^ and spring constant of 1000 kJ mol^-1^ nm^-2^. Configurations were extracted from the pulling simulations at 0.1 nm intervals until the C5aR-PMX53 COM-COM distance was 3.0 Å, and at 0.2 nm intervals thereafter until the final COM-COM distance was 6.0 Å. In total, 36 configurations were generated to serve as umbrella sampling windows. Each of the 36 umbrella sampling windows were equilibrated for 100 ps in an NPT ensemble followed by a 40 ns production run while applying a 1000 kJ mol^-1^ nm ^-2^ force constant along the *z*-axis on the PMX53 molecule. Finally, the free energy profile of transferring PMX53 from its bound state to an unbound state was calculated using the weighted histogram analysis (WHAM) as implemented in GROMACS v5.1.2. Bootstrap analysis was used for estimation of statistical errors.

### Analysis of non-covalent interactions

The various non-covalent interactions were estimated using built-in GROMACS tools and in-house Perl scripts. Hydrogen bonds were estimated using the gmx hbond tool using default criteria. Cation-π and salt-bridge interactions were defined based on the distance criteria described elsewhere.

## Author contributions

A.C.D. and M.K. set up and performed the AFM experiments. D.A. and A.C.D. coanalyzed the experimental and performed calculations. C.Z. and H.L. provided some of the ligands and cloned, purified and reconstituted C5aR. H.F. and R.N.V.K.D. set up and performed the MD/SMD simulations and analysis. All authors wrote the paper.

## Acknowledgements

This work was supported by the Fonds National de la Recherche Scientifique (F.R.S.-FNRS grant numbers: PDR T.0090.15 to D. Alsteens), the Research Department of the Communauté française de Belgique (Concerted Research Action), the Université catholique de Louvain (Fonds Spéciaux de Recherche), the ‘MOVE-IN Louvain’ Incoming post-doc Fellowship programme to A. Dumitru and the National Institute of Health of United States (1R35GM128641-01 to C. Zhang.). D. Alsteens. is Research Associate at the FNRS. H. Fan gratefully acknowledges financial support from Biomedical Research Council of A*STAR. This work was also supported by funding from National Institute of Health. The computational work was performed on resources of the National Supercomputing Centre, Singapore (https:///www.nscc.sg). The authors thank M. Mathelie-Guinlet (Université catholique de Louvain) and A. Vilquin (École supérieure de physique et de chimie industrielles de la ville de Paris) for the support with MATLAB routine coding.

